# Factors influencing the accuracy and precision in dating single gene trees

**DOI:** 10.1101/2020.08.24.264671

**Authors:** Guillaume Louvel, Hugues Roest Crollius

**Author notes:** Corresponding author: Guillaume Louvel.

## Abstract

Molecular dating is the inference of divergence time from genetic sequences. Knowing the time of appearance of a taxon sets the evolutionary context by connecting it with past ecosystems and species. Knowing the divergence times of gene lineages would provide a context to understand adaptation at the genomic level. However, molecular clock inference faces uncertainty due to the variability of the rate of substitution between species, between genes and between sites within genes. When dating speciations, per-lineage rate variability can be informed by fossil calibrations, and gene-specific rates can be either averaged out or modeled by concatenating multiple genes. By contrast when dating gene-specific events, fossil calibrations only inform about speciation nodes and concatenation does not apply to divergences other than speciations.

This study aims at benchmarking the accuracy of molecular dating applied to single gene trees, and identify how it is affected by gene tree characteristics. We analyze 5205 alignments of genes from 21 Primates in which no duplication or loss is observed. We also simulated alignments based on characteristics from Primates under a relaxed clock model, to analyze the dating accuracy. Divergence times were estimated with the bayesian program Beast2.

From the empirical dataset, we find that the date estimates deviate more from the median age with shorter alignments, high rate heterogeneity between branches and low average rate, features that underlie the amount of dating information in alignments, hence statistical power. The smallest deviation is associated with core biological functions such as ATP binding, cellular organization and anatomical development, categories that are expected to be under strong negative selection.We then investigated the accuracy of dating with simulated alignments, by controlling the three above parameters separately. It confirmed the factors of precision, but also revealed biases when branch rates are highly heterogeneous. This suggests that in the case of the relaxed uncorrelated molecular clock, biases arise from the tree prior when calibrations are lacking and rate heterogeneity is high. Our study finally reports the scale of the gene tree features that influence the dating consistency with median ages, so that comparisons can be made with other genes and taxa. To tackle the molecular dating of events only observed in single gene trees, like deep coalescence, horizontal gene transfers and gene duplications, future models should overcome the lack of power due to limited information from single genes.

## Introduction

The phylogenetic tree is the most prevalent concept for representing evolutionary relationships, but a species tree and a gene tree do not necessarily tell the same story: while nodes in a species tree represent speciations, meaning the divergence of sister species from their parent, nodes in a gene tree represent the divergence of genes which do not result directly from speciation, but from the coalescence of alleles or from locus duplication and horizontal transfer. Therefore reconstructing a species tree generally requires sequences assumed to be orthologous and congruent with the species history; subsequent inferences such as dating taxon divergences usually lump together many genes to increase statistical power, for example by concatenating orthologs. However, the specificities of gene evolution are not only a nuisance that hinders species tree inference, they are also the raw material of morphological evolution, adaptation and species diversification. In this regard, focusing on single genes instead of concatenations can reveal gene-specific evolutionary processes. In particular, genes have heterogeneous substitution dynamics due to functional specificities or location in the genome, or as a consequence of events of duplication and transfer. Estimating the rate and timing of sequence evolution is the subject of molecular clock dating and we evaluate here its accuracy and precision when applied to single genes.

The molecular clock is a set of methods to date the divergence between DNA or protein sequences by linking absolute time to the number of observed substitutions (Zuckerkandl & Pauling, 1962, 1965; Margoliash, 1963; Doolittle & Blombäck, 1964). The model considers a sequence that evolves by substitution over a given lapse of time *t.* It states an approximately proportional relationship between time *t* and the number of substitutions *N*, by modelling *N* as a Poisson distributed random variable of expectation *r×t*, where *r* is the rate of substitutions per unit of time. Thus an average of *r*×*t* substitutions occur in the duration *t*. In the simplest model the rate is assumed to be constant over the entire phylogenetic tree (Langley & Fitch, 1974). The number of substitutions can be obtained from the branch lengths inferred with classic phylogenetic tree inference programs (PhyML, Iqtree, RaxML, Phylobayes, MrBayes, PAUP, etc), but to infer the rate, times must be known, and vice-versa. Fossils typically provide independent dating and are used to assign ages to some of the speciation nodes, allowing the rate to be derived and all non-calibrated nodes to be dated.

However while rate constancy is a convenient simplification, it has been observed that rates of substitution per unit of time vary across taxa (Wu & Li, 1985; Britten, 1986; Pagel et al., 2006). In particular, when considering a single site in an alignment, the across branch rate variation is called heterotachy (literally “different rates”; Philippe et al., 2003). The causes of variations are diverse and may include population size, impact of selection at the molecular level, generation time, efficiency of the DNA repair system, metabolism, etc (Gillespie, 1991). For some of these reasons, rates also vary across the genome (Wolfe et al., 1989). At the scale of a single sequence, the heterogeneity of the rate across branches does not necessarily follow the same pattern between sites, meaning that different sites accelerate or decelerate in an independent manner. This results in a complex pattern of variation across sites and genes and across branches. Some authors (Kumar & Hedges, 1998) have argued that by selecting many genes these variations are attenuated, their mean rate is approximately constant and the molecular clock can still be used, especially in recent clades. Elsewhere it is generally accepted that such simple models can generate overconfident but wrong results (Revell et al., 2005; Wertheim et al., 2010; Duchêne et al., 2015; Lozano-Fernandez et al., 2017).

Initially, tests of constancy such as the relative rate tests (e.g. in Sarich & Wilson, 1966; Fitch & Langley, 1976; Gu & Li, 1992; Tajima, 1993) or the likelihood-ratio test of the clock (Felsenstein, 1981) were developed to assess these rate variations. Nowadays, most models accommodate a non constant molecular clock providing that enough calibrations are specified. How to relax the clock constancy is not settled: autocorrelated rates assume a progressive change between connected branches (Thorne et al., 1998; Kishino et al., 2001) while uncorrelated clock rates just follow an overall distribution across branches (Drummond et al., 2006). In the uncorrelated case, the branch-wise rates are most commonly modeled as a lognormal, gamma or exponential distribution, and the branch lengths in units of substitution are usually obtained by multiplying these rates by the absolute time difference. As a consequence the variance of the branch length scales quadratically with the time difference. Alternatively the “white noise” model is totally uncorrelated at all times (i.e. within branches) and has the interesting property that the variance of the branch length scales only linearly with the time difference (Lepage et al., 2007). Different empirical studies support either the uncorrelated rates model (Drummond et al., 2006; Rannala & Yang, 2007; Linder et al., 2011; Heath et al., 2012) or the autocorrelated clock rates (Thorne et al., 1998; Lepage et al., 2007; dos Reis et al., 2018; Smith et al., 2018), while a mixture of the two models over different timescales has also been proposed (Lartillot et al., 2016; Bletsa et al., 2019). However it is uncertain that autocorrelation can actually be detected (Linder et al., 2011; Ho et al., 2015; Tao et al., 2019) and the modalities of the rate relaxation are likely specific to each taxonomic group, timescale studied and calibration scheme. Algorithmic implementations include widely used software such as Beast 2 (Bouckaert et al., 2019) and MCMCTree (Rannala & Yang, 2007) which exploit the Bayesian framework through Markov Chain Monte Carlo (MCMC) sampling, while some faster alternatives take advantage of least squares strategies, like LSD (To et al., 2016) and treedater (Volz & Frost, 2017). Accelerating probabilistic algorithm has also been made possible by using approximate likelihoods (Thorne et al., 1998; Guindon et al., 2010; dos Reis & Yang, 2011).

In contrast to the early ages of molecular biology in the nineteen-sixties where analyses could only be performed on a single protein at a time such as haemoglobins, cytochrome c or fibrinopeptides (Margoliash, 1963; Doolittle & Blombäck, 1964; Zuckerkandl & Pauling, 1965), modern high-throughput sequencing gives access to hundreds or thousands of genes per genome, providing a more general insight. The modern procedure to date species divergences therefore uses many loci whose sequences are concatenated. This concatenation approach aims to increase the amount of information, thus the precision (Bromham et al., 2000; Lanfear et al., 2010; Smith et al., 2018; Bletsa et al., 2019). Even in a favorable situation with many genes, the accuracy of molecular dating has been questioned, because of flawed procedures such as reusing inferred dates as secondary calibrations without propagating uncertainty (Graur & Martin, 2004; Schenk, 2016), but also because of inherent limitations of the problem (Pulquério & Nichols, 2007; Burbrink & Pyron, 2008; dos Reis & Yang, 2013; Zhu et al., 2015; Kumar & Hedges, 2016; Warnock et al., 2017); the most critical limitations are the difficulty of characterizing the type of clock relaxation and the uncertainty in calibration points themselves (reviewed in Ho, 2014; dos Reis et al., 2015; Mello & Schrago, 2024). Assessing the adequacy of different models of rate variation has been investigated by a number of simulation-based studies (Aris-Brosou & Yang, 2002; Ho et al., 2005; Rannala & Yang, 2007; Battistuzzi et al., 2010; dos Reis et al., 2014; Duchêne et al., 2015) but with a focus on multiple-loci datasets and speciation dates.

If accurate dating requires sufficient sequence information as well as calibrations, this means that inferring dates in single gene or transposable element evolutionary histories is even more difficult than with concatenations of hundreds of genes. Indeed, on such short sequences, low statistical power amplifies the following known limitations of dating by molecular clock: methodological errors in tree topologies and substitution rates, rate variation of multiple biological origins and inherent stochasticity in the substitution process, all of which create a disproportionate amount of noise.

Here we assess molecular clock dating from single gene trees taken individually, and what characteristics of a gene tree are related to precision and accuracy. We first evaluate a dataset of primate genes under a cross-validation procedure where we date speciation events in each tree and compare each individual tree estimate with the median age over all trees. Using the median age as a point of reference produces a measure of the precision of dating. In these primate gene trees we find characteristics correlated with the deviation from the median, mainly the length of the sequence alignment, the heterogeneity of the rate between branches and the mean rate of the tree. In addition to measuring the deviation from the median we also compare with reference ages from TimeTree, and observe a bias towards younger ages. Because several explanations for such bias cannot be disentangled from the empirical dataset we simulated gene trees to also measure how the accuracy of dating vary depending on alignment length, rate heterogeneity between branches and mean rate of substitution.

## Results

### Single gene estimates show a high dispersion and a bias towards younger ages

We perform a benchmark on dating speciation nodes from single gene trees with an empirical dataset of trees that include genes from 21 primate species obtained from Ensembl version 93. We focus on the Simiiformes clade as ingroup, which contains all Primates but the Lemuriformes that we use as outgroup. Speciation ages for all nodes have been estimated many times independently based on fossil-calibrated dating on concatenation of genes, and resulting consensus ages can be obtained from TimeTree (Kumar et al., 2017). Because our aim is to replicate the uncertainty in dating nodes that lack fossil calibrations, we do not set any calibration except for the Simiiformes ancestor at 43.2 My (C.I [41.0, 45.7]). We then quantify the uncertainty relative to the surrounding interval of 43.2 My. The choice of this root calibration and its associated uncertainty is arbitrary because all trees are then compared by this yardstick. Likewise, the choice of Primates for the source trees is arbitrary; the specific selection of species does not matter, what matters is that we collect natural replicates of the same tree. We selected 5205 gene trees that do not display any duplication or loss in this tree of 21 primate species, so that each gene tree shares exactly the same topology. These were dated with Beast 2, providing 5205 age estimates at each internal node (fig. 1). Because the gene trees were independently subjected to dating, and the dated output trees were all scaled to 43.2 My, their differences in mean rate (i.e. the “gene effect”, Ho 2014) are not considered in this part. The impact of genes specific features is investigated in the subsequent part.

**Figure 1:**
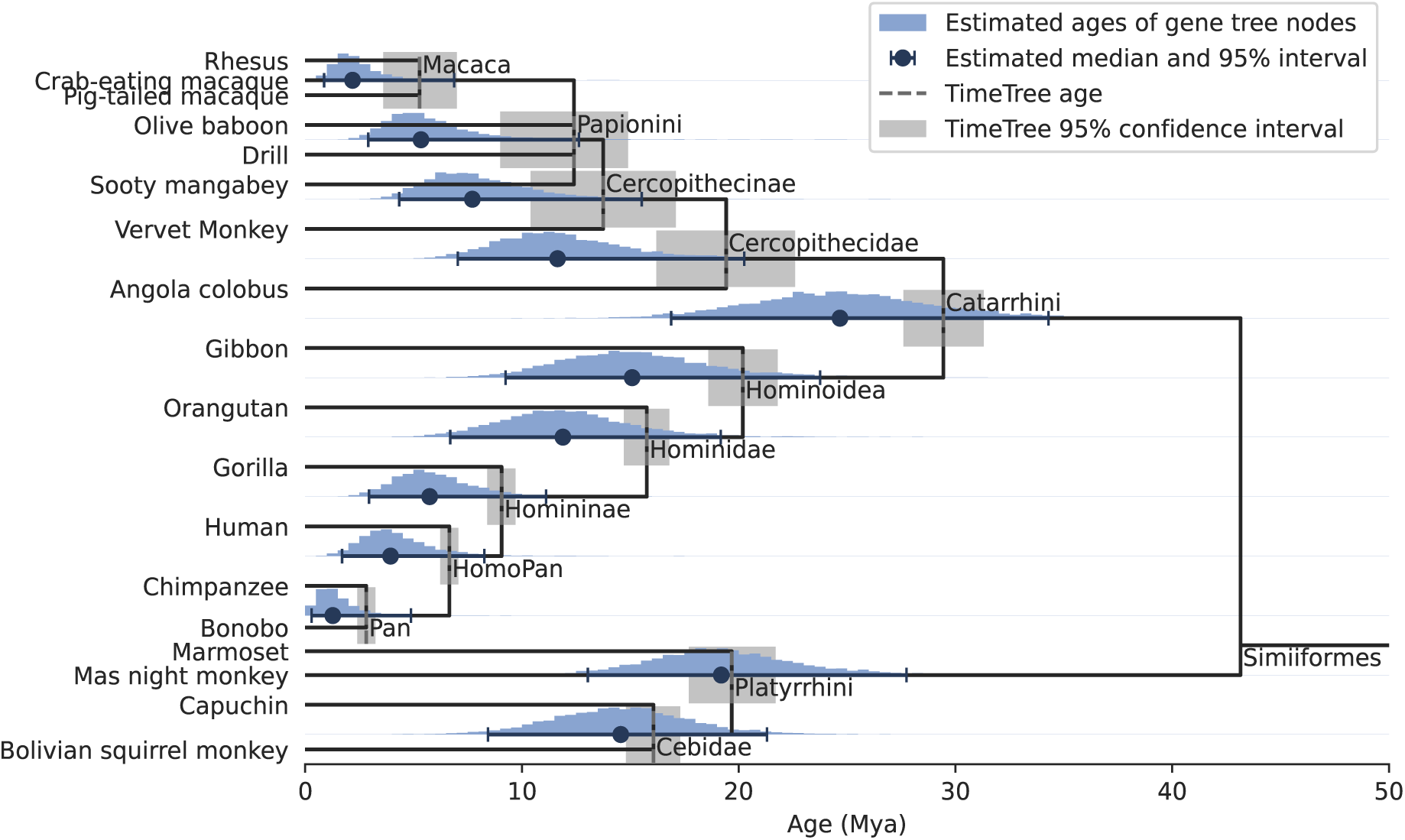
Distribution of speciation ages estimated on primate gene trees independently (5205 trees). Each histogram is rescaled to display the same height, but their total count is each equal to 5205. Dating was performed with Beast under the nucleotide substitution model HKY and a relaxed clock rate (see Methods).

By comparison with the reference age estimates from TimeTree, we obtain clearly younger estimates on all nodes, sometimes even outside of the confidence interval from the reference (median, fig. 1). The highest shift of 10 My affects Cercopithecidae. This suggests that either our estimates are biased, or less plausibly that the reference ages themselves are inaccurate. In our dating procedure, several simplifications may bias ages. To start with, calibrating only one node is unlikely to lead to accurate variable rates, but this is precisely the purpose of our analysis, since we study gene trees for which nodes lack calibrations. With one calibration but variable branch rate, rate and time are unknown because they are conflated into the branch length. In fact, molecular clock “dating” is as much about estimating rates using calibrations than estimating dates from rates, this shift in focus being due to the high variability of molecular rates. Without many calibrations, variation in rates cannot be faithfully inferred, and in turn dates remain uncertain. Accordingly, our empirical dataset is expected to display heterogeneous branch rates of different origins. First there are species-specific rate variations, in which all genes experience the same trend in a specific lineage (also called lineage effects; Gillespie 1989; Muse and Gaut 1997; Smith and Eyre-Walker 2003). For example the average branch length of Cercopithecidae is higher than their Hominoidea sister (supp. info. S1). This is consistent with the known generation times in these clades: based on Pacifici et al. (2013) the averages are 4035 days for Cercopithecidae versus 6132 for Hominoidea. In addition to species trends, each gene tree may experience particular variations of branch rates, therefore producing dispersed ages when compared (called residual effects, i.e. variation remaining after gene and lineage effects; Gillespie 1991). Finally, independent across-branch rate variation also likely occurs within genes, between sites. However, these gene and site heterotachies should just cause dispersion but not loss of accuracy, as we expect that the distortions on branch lengths should compensate themselves on average (after correction for lineage specific rates). Another simplification of our inference is to consider instantaneous segregation of genes at speciations. In reality, speciation takes up to several million years and segregated genes can correspond to older allelic divergence. This phenomenon of deep coalescence should cause speciations to appear older than they are, and not younger as is the case in our results. Conversely, shallow coalescence caused by introgression between recently diverged species would produce younger age, but we deem implausible that introgression would be so pervasive among genes so as to strongly bias ages. In summary, variation in lineage specific rates is the most plausible explanation for our bias towards younger ages, unless the inference obtained from Beast is not statistically consistent.

Aside from comparisons with the reference ages, we quantified the dispersion of the estimates at each node with the following metrics: the mean absolute deviation from the median (MAD) of the node age and the 95% inter-quantile range (IQR95); the averages of these metrics over the 12 speciations yield 3 My and 16 My, respectively. These dispersions are larger for deeper nodes, such as Catarrhini with a MAD of 5.6 My and an IQR95 of 32.4 My. Divided by the Simiiformes calibration of 43.2 My, they represent 13 % and 75 % of this interval, respectively. In other words, the confidence interval for the age of Catarrhini is barely shorter than the age of Simiiformes.

In the following, we break down the causes leading to the dispersion of our estimates by measuring gene features and searching for a relationship with the deviation from the median ages.

### The dispersion of age estimates is associated with low statistical signal and branch rate variation

We computed 56 features from each gene tree related to alignment quality and substitution rate estimations (supp. info. S2). We regressed them against the absolute deviation from the median age averaged across all speciation nodes, chosen to describe the dating imprecision in a single gene tree. In order to select a restricted set of independent variables we used a Lasso regression with a regularization parameter ‘alpha’ of 0.02, which retained 10 parameters further fitted by Ordinary Least Squares (adjusted *R²* = 0.288, fig. 2). Variables were normalized so that coefficients can be compared and they were centered. Apparent outliers, such as trees with extreme rates, were discarded from the regression (see Methods - Removing trees with outlier values), which was therefore based on 5170 trees.

**Figure 2:**
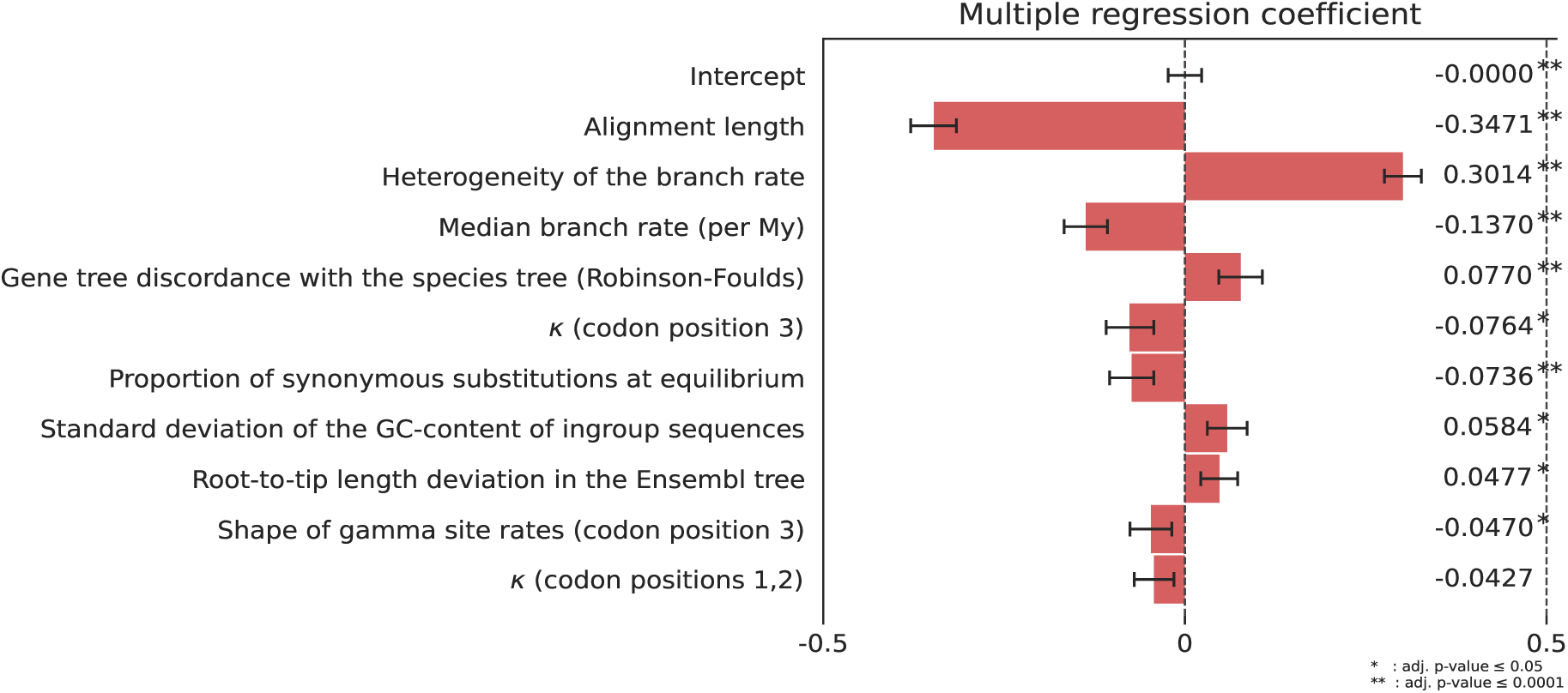
Coefficients of regression of the gene tree features against the age dispersion. Features were selected with a Lasso regression with parameter alpha=0.02. Adjusted *R²* = 0.288. Error bars indicate the 95% confidence intervals.

By order of coefficient size, the length of the sequence alignment comes first (coefficient 0.35), associating short alignments with high dispersion. It is followed by the rate heterogeneity across branches which is positively associated with the dispersion (0.30). Next comes the mean rate of substitution (−0.14). We hypothesize two distinct causes for these correlations: first, the strength of the statistical signal is linked to the alignment length and to the mean rate, because both influence the number of observable substitutions, thus the amount of information available to infer ages; also because of rescaling all trees to the same height of 43.2 My, trees with the lower mean rate are expected to display a higher dispersion. Second, some model parameters are in practice difficult to infer individually. In particular, a variability in branch rates cannot be estimated when there is only one calibration, meaning that a branch specific rate could take arbitrary values as long as the product of rate by time equals the branch length in substitutions. Consequently, it would take values dictated by the tree prior (the Birth-Death branch process) that cannot possibly recover the exact Simiiformes speciation dates (dos Reis & Yang, 2013). Here the rate heterogeneity is the standard deviation of the rate across branches, as inferred by Beast. This indicates that while Beast manages to detect a high rate heterogeneity in a given gene tree, the ages it infers in such a tree are quite distinct from the median ages, and probably far from their true age. The fourth largest association is a measure of the incongruence between the species tree and the gene tree reestimated using IQtree. This means that lack of support for the species topology is associated with high deviation from the median ages. This could be attributable to either actual incongruence being masked by the reconciliation step in our dataset, to low signal from short sequences or to such departure from a strict clock that the true topology cannot be recovered.

The next five coefficients have p-values below 0.05, and correspond to characteristics of the substitution process or the alignment, for which we provide the following interpretations. The kappa parameter (for codon position 3), or ratio of transitions over transversions, is negatively correlated. It may relate to the degree of saturation: saturation brings this ratio closer to one, and it should increase dating uncertainty, although we think that very few gene trees show saturation in our Primates dataset. Next is the proportion of synonymous substitutions at equilibrium, negatively correlated. This value that we measured with ‘codeml’ from PAML depends on the alignment and represents the amount of substitutions that would be synonymous if the ratio dN/dS was equal to one. This result indicates that gene sequences that offer more opportunity for synonymous changes favor precise dating. For the standard deviation of the GC-content between sequences of the alignment, the correlation is positive. This could have two causes that are not mutually exclusive, one methodological, one biological: in the substitution model, sequence composition is assumed homogeneous. Only few phylogenetic models can actually model composition changes over the tree (Foster, 2004) while this is known to highly impede phylogenetic inferences. Under a purely biological explanation so even if we fitted the appropriate model, genes subject to composition shifts could be evolving heterogeneously in other aspects, in particular with heterogeneous rates. The next variable is the standard deviation of root-to-tip path lengths, an approximation of the branch rate heterogeneity that can easily be computed on any phylogram. It is, as expected negatively correlated but weakly, probably because our more precise measure of across-branch rate heterogeneity already captures most of the effect. In Smith et al. (2018) it is used as a criterion for selecting trees which are suitable for dating, and we can expect a better predictive power by using a more refined measure of rate heterogeneity instead; we also performed the regression with only the root-to-tip variance as the sole measure of rate heterogeneity, and it has a lower coefficient than the more refined measure (supp. info. S3). Last to be significant, the shape of the site-wise gamma distribution of rates is a parameter in which a lower value means a more skewed gamma distribution. Thus, as indicated by the negative coefficient (fig. 2), rates that are heterogeneous site-wise (not only branch-wise) appear to also increase dating uncertainty.

To situate these features in their real scale, we summarized them in the 10% of trees with highest or lowest predicted dispersion (table 1). Additionally, we searched for gene functions overrepresented in these two extreme subsets compared to the full set of 5170 genes, based on human Gene Ontology (GO) annotations (table 2, supp. info. S4). For the most dispersed trees no enriched function was detected. On the other hand, well dated trees (low dispersion) appear to have constrained functions. Negative selection on this set is further evidenced by 67 “Human Phenotype Ontology” overrepresented terms related to various developmental abnormalities (nervous system, eye, ear morphology, movement, see supp. info. S4). These functions are a subset of the functions retrieved when using the 10% longest genes (supp. info. S5), indicating that the length is the main driver of the functions found in the set of genes that have the lowest dating deviation from the median.

**Table 1:** Characteristics of gene families whose dispersion is predicted to be the lowest/highest (mean ± standard deviation).

| Subset of gene families | Observed dispersion (My) | Alignment length (nucl) | Rate heterogeneity ($10^{-4}$ subst/nucl/My) | Mean rate ($10^{-3}$ subst/nucl/My) |
| --- | --- | --- | --- | --- |
| 10% highest predicted dispersion | $3.45 \pm 1.6$ | $1008 \pm 506$ | $11.3 \pm 9.76$ | $2.16 \pm 1.25$ |
| 10% lowest predicted dispersion | $1.41 \pm 0.576$ | $5255 \pm 3247$ | $4.69 \pm 3.94$ | $2.75 \pm 1.13$ |

**Table 2:** Overrepresented Gene Ontology terms in gene families with lowest/highest predicted dispersion, using g:Profiler with default parameters: significance threshold 0.05 and multiple testing correction g:SCS.

|  | GO category | GO term IDs | GO term names |
| --- | --- | --- | --- |
| 10% lowest predicted dispersion | Molecular Function | GO:0005524 | ATP binding |
|  |  | GO:0032559 | Adenyl ribonucleotide binding |
|  |  | GO:0030554 | Adenyl nucleotide binding |
|  | Biological Process | GO:0071840 | Cellular component organization or biogenesis |
|  |  | GO:0016043 | Cellular component organization |
|  |  | GO:0007010 | Cytoskeleton organization |
|  |  | GO:0006996 | Organelle organization |
|  | Cellular Compartment | GO:0005604 | Basement membrane |
|  |  | GO:0043228 | Non-membrane-bounded organelle |
|  |  | GO:0043232 | intracellular non-membrane-bounded organelle |

In this regression of gene tree characteristics, the explained variable (deviation from the median) is a measure of dispersion, not of accuracy. However we can only measure accuracy if we have access to the “true” age. Besides, the distribution of ages (fig. 1) suggests a bias in our estimates that empirical data alone is powerless to explain. Furthermore, empirical data is the product of several confounding factors, where for example an apparent rate heterogeneity might instead be caused by undetected incongruence in the gene tree. We therefore performed a simulation to confirm the features that impact dating accuracy.

### Simulating gene tree variability impact on dating

Simulations allow to measure two facets of the dating quality: first the precision or how fine-grained the estimates are; second and more importantly, the accuracy or how close the estimates are from the true value on average. We focused on the top 3 variables obtained from the regression: alignment length, across-branch rate heterogeneity and mean rate of substitution. Our three variables of interest were set to representative values spanning their observed range in the real trees (supp. info. S6). For each set of parameters, we first simulated branch lengths on the fixed primate tree according to a relaxed clock model, then simulated codon sequence evolution along this tree (see Methods - Simulating alignments). Dates were then reconstructed from simulated data with the same method as for the empirical dataset. We expected the three variables to reproduce the same effect on dispersion as with the above empirical dataset. However we expected the accuracy to be unaffected because the model we use to simulate alignments and trees is very similar to the model used to reconstruct dates: Beast dating was thus performed without discrepancy between its inference model and the process that generated the data, and in this condition we expect it to be statistically consistent, i.e. converge to the true value as the amount of data increases.

We ran 500 simulations for each parametrization, by fixing two parameters and varying the third, around a central point defined by length=3000 nucleotides, mean rate=0.004 substitutions/codon/My and branch rate heterogeneity=¼ (defined as the standard deviation divided by the mean of the branch rates, abbreviated *σ/µ*).

The dispersion, as seen by the interquantile range covering 95% of the data (IQR95) (fig. 3) varies with each variable in the same direction as observed in the regression of primate gene trees: it increases for shorter alignments (3b), higher rate heterogeneity (3c) and lower evolutionary rate (3d). In almost all speciation nodes and parameter values, we recover an unbiased median age, falling accurately on the age from the underlying species tree. However, we find shifts for the highest rate heterogeneity between branches (3d, *σ/µ* = 1). Cebidae appears younger while Catarrhini appears older than in reality, an effect that we find when sampling from the prior (supp. info. S7). This shows that in presence of very high across-branch rate variation and uninformative calibrations, the prior on the time tree (Birth-Death) strongly influences the ages.

**Figure 3:**
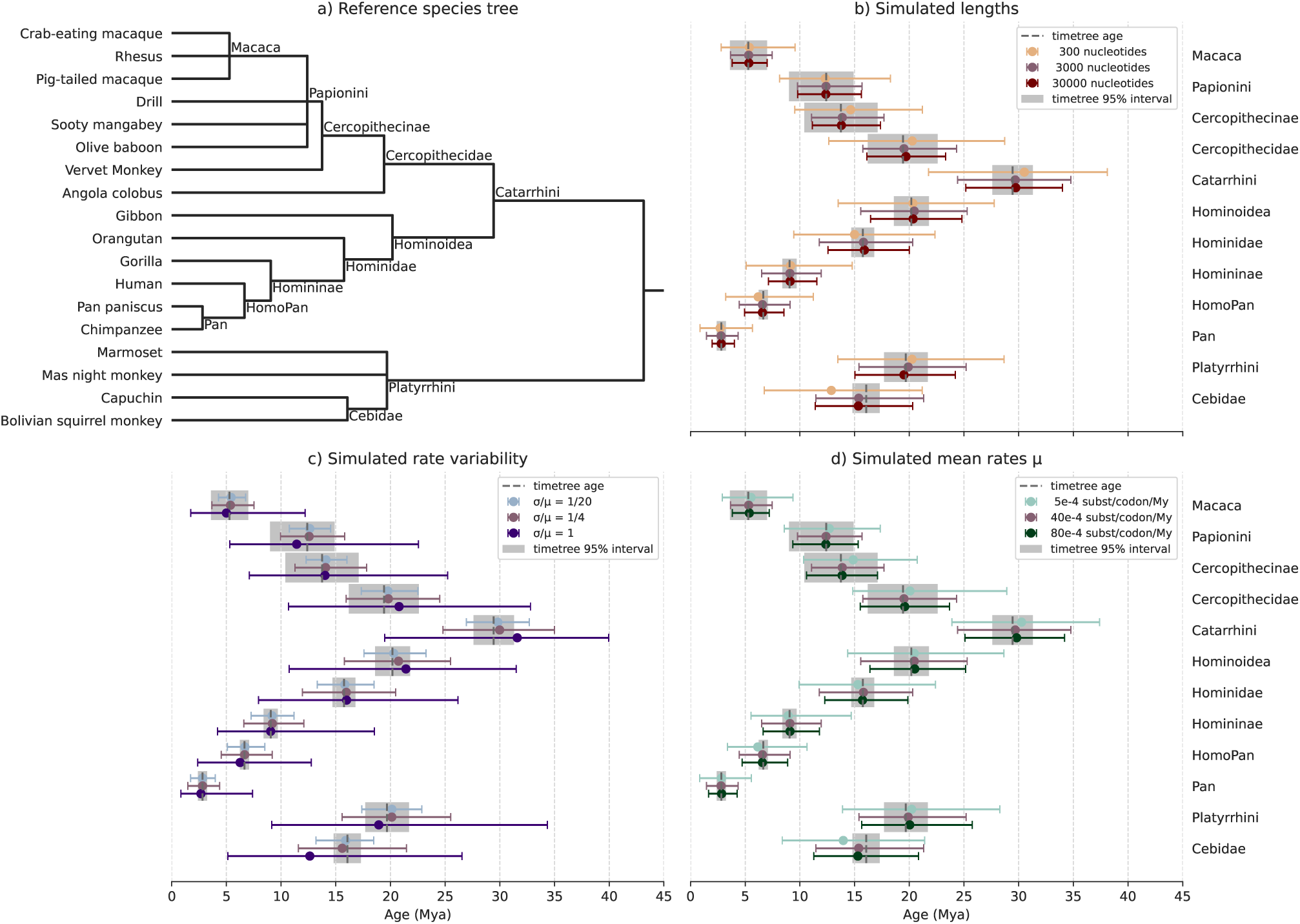
Speciation dates on simulated alignments, under a relaxed clock model and a fixed species tree of Simiiformes, under various parameter values. Central dots represent the median date, and the interval represent the range [0.025, 0.975] of the distribution. a) Dated species tree from TimeTree; b) the number of sites in the alignments, in number of nucleotides; c) the diffusion parameter, defining the degree of relaxation of the clock rate, such that the ratio *σ/µ* (standard deviation of the rates over the mean rate) equals one twentieth, one quarter, and one; d) The mean rate of substitution, in number of substitutions per site per million years. When not specified, length is 3000, rate is 40e-4 and *σ/µ* is ¼.

## Discussion

### Factors of consistency between ages inferred from separate gene trees

From an empirical dataset of 5205 gene trees in 21 Primates, we identified characteristics related to the deviation of age estimates from the median age. More statistical power, provided by higher rates or longer alignments, lowers the dating deviation while an increase in rate variation between branches increases the deviation. This confirms previous observations that the number of sites and substitutions is a limiting factor with regard to statistical power (Bromham et al., 2000; Lanfear et al., 2010; Smith et al., 2018; Bletsa et al., 2019; Bromham, 2019), and also that the clock rate does not appear to be constant across branches. In our analysis the deviation also correlates with characteristics of the substitution process (kappa, GC content), indicating either that the dating model is too simple, or that multiple kinds of heterogeneities (composition, rates) co-occur in some trees. The 5205 selected gene trees only contain strict orthologs and as such represent a biased fraction of the 24,614 gene trees extracted from Simiiformes: this selection likely represents a more reliable subset for dating, because the absence of gene duplication and loss may be associated with a more stable evolution in terms of constancy of rate and selective pressure. On the other hand, even this selection of trees can contain incongruent trees not identified as such, because of the reconciliation method that forces gene trees into the species topology without a model for incomplete lineage sorting (ILS) or introgression. This is suggested by the fourth detected factor in our regression being the discordance between the recomputed gene trees and the species tree. Despite having kept a few polytomic species nodes to lift some of the constraints on the topology, it is likely that it still distorts branch lengths (Mendes & Hahn, 2016; Carruthers et al., 2022).

In the regression based on empirical gene trees, the adjusted *R²* is low (0.29), indicating a low fraction of explained variance. To investigate the missing variance, we should incorporate additional biological characteristics of the sequences, but as we show in simulations, stochastic properties of the mutational and evolutionary processes already account for a large uncertainty. Furthermore in our empirical dataset, ILS, introgression or reciprocal paralog loss may also obfuscate the relationship between molecular change and time, but we have not used models able to identify these processes. Among other simplifications we used, our dating method employs a nucleotide substitution model which cannot distinguish neutral from non-neutral substitutions. According to the nearly neutral theory the majority of amino-acid changes is slightly deleterious which causes molecular divergence patterns to be more clock-like in absolute time, whereas strictly neutral substitutions should show a generation time effect (Ho, 2014). A codon model would be able to separate synonymous from non-synonymous substitutions and test this hypothesis, but being more complex (using 61 character frequencies instead of 4) it also requires more data, which as we have seen is the limiting factor at the scale of single gene trees.

### Limiting the bias when dating single gene trees

In the empirical estimates, we observed a shift towards ages younger than the TimeTree reference. Then we simulated sequences without species-specific rates and that bias was not reproduced. This is unsurprising because we used the same model for inference as for simulating the data, and thus were expecting statistical consistency. However in the simulated case with very high rate heterogeneity, the ages are biased by the time tree prior (Birth-Death model). Other sources of bias have been demonstrated, in particular incongruence with the species tree as caused by incomplete lineage transfer (ILS) (Mendes & Hahn, 2016; Carruthers et al., 2022). In the latter extensive empirical and simulation study, it was shown that the length of branches descending from incongruent nodes (such as terminal branches) was overestimated, while the length of branches predating incongruent nodes was underestimated. This produces older ages than the actual speciation times, and the authors propose to mitigate this by considering congruent branches only. In the real primate trees we observe younger ages instead so it is likely that the bias is instead driven by species-specific rates, a factor that was absent from our simulations. Other studies on empirical data have found biases caused by heterotachy (Wertheim et al., 2012), especially if the rate change is punctuated (Dornburg et al., 2012). In presence of substantial lineage rate variation the relaxed clock model requires calibrations on internal nodes to properly infer branch rates and times (Duchêne et al., 2014). Here, the choice of a single calibration was made precisely with the aim of measuring the dating accuracy on uncalibrated nodes, such as those occurring in gene trees if we consider events other than speciations. Some studies use the relative ordering of events from different taxa (Lutzoni et al., 2018) or from different gene families (Pittis & Gabaldón, 2016; Vosseberg et al., 2021), but across-branch rate heterogeneity is a major hurdle to doing so and conditions for applicability need to be rigorously verified, in particular the absence of systematic bias (Susko et al., 2021).

### How to reduce uncertainty

*Stochastic uncertainty* is the amount of information that is lost assuming that the model is correct, as opposed to *inductive uncertainty* due to inadequate modelling (Holland, 2013). Inadequate modelling generally produces systematic errors. Our simulations show what level of stochastic uncertainty to expect in gene trees with realistic rate characteristics, and they confirm the drivers of uncertainty identified in empirical gene trees. We highlight the requirement for a high number of sites and substitutions, which is critical for statistical power. This problem can probably only be circumvented by incorporating information beyond single genes, such as genomic context and linked model parameters between multiple gene trees, as recent developments suggest (Duchêne et al., 2020). However fossil calibrations are quantitatively as important as sequence data alone to accurate dating (dos Reis et al., 2015). These solutions are unfortunately harder to design for gene trees with duplications or transfers because they do not share the same set of branches. Some pragmatic approaches that have been proposed to handle gene specific heterogeneity are the removal of loci with apparently non constant rate prior to the analysis (Jarvis et al., 2014), a “gene-shopping” approach (Smith et al., 2018), but discarding data might be unsatisfactory for genome scale analyses that look for the most general picture. Our results may be extended to a similar gene filtering approach as it pinpoints gene features that are correlated with the dating uncertainty, although its predictive power would need to be improved. Beyond identifying outliers, it would be interesting to understand why they are so, in terms of function, selection pressure or genomic context. In this high-throughput era, homologous sequences from a wide variety of taxa are accessible. A few genes have received considerable attention because of very distinctive evolutionary dynamics, such as PRDM9 which is generally evolving under positive selection and is a “speciation gene” in mammals (Oliver et al., 2009) or MHC immune genes that display elevated polymorphism in populations (Piertney & Oliver, 2006). Since our evaluation used only one calibration, it is a worst case setting that could occur in gene trees with many duplications or transfers, events for which fossil calibration is less informative (but see Davín et al., 2018, who use horizontal transfers as relative time constraints). Key adaptations or transitions may result from these gene specific events, in particular duplications (Aguileta et al., 2006; Vosseberg et al., 2021), horizontal and endosymbiotic transfers of genes (Ochman et al., 2000; Koonin, 2016), or movements of transposable elements (Boissinot et al., 2000; Ovchinnikov et al., 2002; Khan et al., 2006).

Regarding *inductive uncertainty*, richer models avoid overconfident but erroneous results. Notably, there is room for refining the models of sequence evolution, for example by considering indels evolution, or the domain composition of proteins, or their 3D structure and the consequence on residue coevolution. Also, when the inference of tree topology, substitutions and clock rates is performed jointly by an integrative method such as Beast, it guarantees that uncertainty is properly cumulated at all steps. However in our experience it appears challenging to run a complex parametric inference such as Beast at the scale of comparative genomics. Such integrative methods might not be numerically tractable because they generally require MCMC which implies prohibitive running times and in addition, MCMC requires a thorough and experimented human inspection to validate the inference output. In the face of this, numerous projects aiming at sequencing broad segments of biodiversity on Earth are emerging, under the auspices of the Earth Biogenome Project (EBP; Lewin et al., 2018, 2022). Extremely large datasets, composed of tens of thousands of genes, will become common place. Therefore the development of faster non-parametric algorithms is still relevant, alongside improvements in the computational footprint of probabilistic algorithms (Mello & Schrago, 2024).

Research on the molecular clock enters an exciting time, with huge amounts of data and increasingly sophisticated methods to dissect the hidden mechanisms of sequence evolution. How the rate of molecular evolution varies still holds mysteries, both at the scale of lineages and at the scale of genomes. Future developments answering this question will be methodologically challenging but shall shed light on important evolutionary processes.

## Methods

### Source and number of trees

The species tree and gene trees were obtained from Ensembl Compara 93 (July 2018; Zerbino et al., 2018) Metazoa dataset: 99 species descending from Opisthokonta (the last common ancestor of fungi and metazoan) and 23,904 reconciled gene trees. Low quality genomes, aberrant gene branch lengths, split genes were removed (supp. info. S8, Species and gene trees preprocessing).

Nodes from the species tree were assigned an age from TimeTree (data retrieved Jan. 2019; Kumar et al., 2017). The corresponding sequences of the coding domain of the longest transcribed isoforms (fasta format) were downloaded as fasta multiple sequence alignments via the Ensembl Perl API.

Focusing on the 21 primate species (18 Simiiformes and 3 Lemuriformes), we extracted 24,614 subtrees at their Simiiformes node plus two of the outgroup sequences with shortest branch lengths.

From those we selected the 5235 gene trees without duplications and losses within Simiiformes, so that they share a single tree topology.

### Multiple Sequence Alignment building and cleaning

We built codon alignments: protein sequences were aligned using FSA version 1.15.9 (Fast Statistical Alignment, Bradley et al., 2009) with default parameters, then back-translated to the corresponding nucleotide (codon) alignment.

HmmCleaner (version 0.180750; Di Franco et al., 2019) finds segments of sequence that appear inconsistent with the other aligned sequences, interpreting them as sequencing errors. We apply it and replace the detected segments by gaps. It was applied to the 5235 alignments that do not have duplications or losses, on the amino-acid data which were then backtranslated to codons. Out of these, 30 alignments caused HmmCleaner to fail, resulting in the 5205 subtrees under study here.

### Bayesian node dating with Beast 2

Using the alignments as input for Beast 2.6.3 (Bouckaert et al., 2019) we ran the following model: a HKY nucleotide substitution model with two site partitions corresponding to the codon positions {1,2} and 3. The “Birth-Death model” is set as the tree prior (Gernhard, 2008), and a relaxed uncorrelated lognormal clock rate is fitted on branches, with mean and standard deviation being estimated. All 12 Simiiformes clades (see species tree in fig. 1) as well as Primates and Lemuriformes were constrained to be monophyletic (note that Macaca, Papionini and Platyrrhini are polytomic). Only Primates and Simiiformes were calibrated precisely, the others having uninformative priors. The calibrations were specified as gamma distributions manually tuned to match TimeTree point estimate and 95% interval: offset 70.8 My, shape 4.6 and scale 0.656 for Primates, and offset 40.9 My, shape 4.0 and scale 0.575 for Simiiformes (therefore the mean prior age for Primates is 70.8 + 4.6×0.656 = 73.8 My, and for Simiiformes it is 40.9 + 4.0×0.575 = 43.2 My). BEASTGen (v1.0.1) was used to automatically generate the parameter file for each alignment from a common template. The MCMC was run in one chain of 20,000,000 iterations, after a pre-burnin of 1,000,000. The 390 trees which had an ESS (effective sample size) under 200 for any variable were resumed and extended with 20,000,000 additional iterations, leading to 13 trees with one ESS under 200 that we discarded. The resulting mean dates were annotated on the species tree using TreeAnnotator from Beast 2.

### Global age dispersion from all primate trees

Given that all speciation nodes in the primate species tree are replicated in our set of 5205 gene trees, we measured the dispersion of estimated ages for each speciation. We used two metrics of dispersion, the MAD and the IQR95. The mean absolute deviation from the median (MAD) is a robust estimate of dispersion: it is less influenced by skewed distributions than when using the mean as center, and less influenced by outliers than the standard deviation. It is also still sensitive enough compared to median deviation from the median or interquartile range. The IQR95, or interquantile range covering 95% of the data (between percentiles 2.5 and 97.5) is more robust but less sensitive.

### Regression

Statistical analyses on the output dates were performed with Python 3.8 and additional scientific computing packages including Numpy, Pandas, Statsmodels, Scikit-learn, Biopython and Ete3 (supp. info. S9).

#### Quantifying the average deviation of ages for a single gene tree

As the dependent variable to regress, we used the average deviation of ages in each gene tree: for one gene tree, the absolute deviation from the median speciation age is obtained at each node, then the deviations of the 12 nodes are averaged.

#### Retrieving mean rate and rate heterogeneity per tree

The clock model that we fit in Beast is unlinked between codon positions {1,2} and position {3}, meaning that the rate parameters are inferred separately for each of these site partitions. Beast outputs different types of rates: First, it outputs the sampled parameters of the uncorrelated log-normal clock model (its mean and standard deviation). Additionally, it computes a posteriori the mean and variance of the branch rate. The latter estimation differs in that it is not a parameter of the model, but a statistic that is computed at the end of each iteration on the proposed tree. We monitor both estimates because they yield quite different values, although being correlated (supp. info. S10). For each sampled tree, it then produces the mean and variance. We use the latter rate statistics as variables in the regression. Since there are as many values as sampled trees of the MCMC chain, we take the median over the chain as final value.

As we estimated separately the mean rates *m*_1,2_ and *m*_3_ for the corresponding codon positions, we finally combined them to obtain the total rate of substitutions per codon:

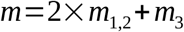

This rate by codon equals three times the average rate by nucleotide, but we chose the codon metric for consistency with the simulation parameters based on a codon model in INDELible.

Similarly as measure of across-branch rate heterogeneity, we summed the rate standard deviations based on the returned variances *v*_1,2_ and *v*_3_ from Beast:

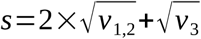

#### Collecting features of the gene families

We measured 56 characteristics of the gene families as explanatory variables in the regression. They include (full list in supp. info. S2):

- Tree features:

– mean bootstrap values of nodes, from Ensembl;

– the root-to-tip deviation, or standard deviation of the path lengths from the root to the leaves, from the Ensembl trees.

– whether the gene tree was a posteriori edited to fit the species tree topology (See supp. info. S8, Species and gene trees preprocessing).

- alignment features:

– global statistics such as alignment length and proportion of pairs of sequences that do not share sites (indicative of “split genes”);

– statistics over sequences like the mean frequency of gaps, the mean percentage of ambiguous nucleotides, the mean GC-content, the mean CpG content;

– statistics over sites, like the column entropy and the parsimony score;

– proportions of sequences cleaned by HmmCleaner;

- substitution process features estimated with the program ‘codeml’ from PAML 4.9e (Yang, 2007):

– proportion of synonymous substitutions at equilibrium, based on measured codon frequencies and the fitted substitution matrix (using names from ‘codeml’ output, it is S/(N+S));

– mean and standard deviation of ω (dN/dS) under the free-ratio model;

- substitution process features estimated during the Beast dating inference, separately for codon positions {1,2} and {3}:

– the ratio of transitions over transversions (kappa);

– the gamma shape for site variable rates;

– the mean clock rate;

– the standard deviation of the clock rate;

- MCMC related statistics of the Beast runs:

– the number of iterations the chain was run (either 20 or 40 millions);

– whether any parameter had an ESS below 200.

#### Transforming regressed features

We chose the appropriate transformation of features in order to stay close to the assumptions of a linear modelling framework with ordinary least squares, that is explanatory variables with low skew. For this, a semi-automated procedure was set up, where the transform that minimizes the skew was selected from the predefined set made of ‘no transform’, ‘logarithm base 10’, and ‘square root’, with additional increment or sign modification if needed by the function domain of validity. Some variables were binary encoded in cases where a large proportion of values was constant, or where distributions were clearly bimodal. Transforms for each variable can be found in supp. info. S11. Finally, all variables were normalized (divided by their standard deviation) and centered before the regressions.

#### Reducing multicollinearity

Multicollinearity being known to impede ordinary least squares fitting, we reduce it prior to the regression with two strategies:

- first we obtained covariances from a Factor Analysis (analogous to Principal Component Analysis, but accounting for discrete and ordinal data). From the results, we removed some features from clusters of heavily correlated features. We also “decorrelated” pairs of features by dividing one by the other or by computing the residues of the simple regression between the two linked variables. For instance we decorrelated standard deviation metrics when they correlate with the mean of the same variable (supp. info. S11).
- Afterwards we checked the multicollinearity condition number, which is the square root of the highest eigenvalue divided by the smallest eigenvalue of *X^T^X* (*X* being the design matrix). Since the multicollinearity condition number was already less than the usual cutoff of 20, it was not needed to remove additional features.

#### Removing trees with outlier values

To avoid a misleading impact of outliers on the linear regression, trees with excessive rates computed by Beast were excluded. The cutoffs were chosen based on the histograms, which display very long tails. As mentionned above, the 13 trees with insufficient ESS (less than 200) were discarded as well. Finally, alignments including pairs of sequences which do not share any common site, i.e. unaligned pairs, were also excluded, as this can lead to meaningless branch lengths. In total these filters removed 35 outlier gene trees (supp. info. S12).

#### Final feature selection with Lasso and OLS refitting

Lasso (Tibshirani, 1996) is a regression fitting algorithm which simultaneously performs feature selection, thus dealing with highly dimensional design matrices. We therefore apply it in a first pass to discard coefficients with an absolute value inferior to 0.01, using a penalty value (parameter alpha) of 0.02. However Lasso coefficients are biased (by design), so that p-values and *R²* cannot be readily computed. For this reason we subsequently refit by ordinary least squares (OLS) using the selected features, with a covariance set to MacKinnon and White’s (1985) heteroscedasticity robust type 1. Finally, the p-values were subjected to a Bonferroni correction by multiplying them by the number of features used in the Lasso step. Statsmodels 0.13.2 implementations were used. Detailed regression output statistics are in supp. info. S13.

### Gene functional annotation overrepresentation

From the 5170 gene trees without duplication or loss retained in the regression, human genes were used as the reference set of functions. Overrepresentation of the 517 human genes from the most dispersed and least dispersed trees was done with the online tool g:Profiler version e111_eg58_p18_30541362 (Raudvere et al., 2019) using default parameters (significance threshold 0.05 and multiple testing correction by g:SCS).

As specified in the table 1, the set of most (or least) dispersed trees is obtained from the fitted regression line. It is therefore a predicted dispersion, distinct from the observed dispersion displayed in column 1.

### Simulating alignments

Alignments were generated by simulating the evolution of a sequence along the above tree of Primates, using INDELible v1.03 (Fletcher & Yang, 2009). The three tested parameters were the mean rate, the rate heterogeneity across branches and the alignment length, with 500 simulation replicates for each set of parameter values. These parameters were chosen to represent the extent of variation in the real data of 5205 trees (as measured to be used in the regression, see below and supp. info. S6). To generate the variable branch rates, we applied an independent log-normal relaxed clock simulation with the ‘simclock’ R package; the tested mean rates were 5×10^-4^, 40×10^-4^ and 80×10^-4^ substitutions.site^-1^.My^-1^. To obtain the desired rate heterogeneities, we tuned the “diffusion” parameter *s²* so that the standard deviation of the branch rates is comparable with empirical estimates, as per the relation between standard deviation *σ*, mean *µ* and diffusion *s²* of a log-normal distribution:

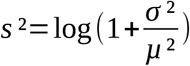

The chosen diffusion values (*s²*) of 0.6931, 0.0606 and 0.0025 correspond to a standard deviation of the branch rate equal to 1, 1/4 and 1/20 of the mean rate (*σ/µ)*, respectively. For the alignment length, we set the root sequence lengths to 300, 3000 or 30,000 nucleotides in INDELible. The evolutionary model includes insertions and deletions, which were parametrized to occur at a rate of 8.4×10^-7^ and 16.8×10^-7^ My^-1^ respectively (corresponding to 1/14 and 2/14 of the median substitution rate 1.177 codon^-1^.My^-1^, a ratio taken from Fletcher and Yang 2009), with lengths following a Lavalette distribution with parameters (2, 300). The sequences evolve according to a codon model with *kappa* = 4 (transitions/transversions) based on estimates from the Primates gene trees (supp. info. S14), *omega* = 0.175 (dN/dS) and codon frequencies taken from the concatenated alignments of Simiiformes genes. No across-site rate heterogeneity was modeled. The random seed was set to 9342.

We then dated Primates speciations from these simulated sequences with Beast 2, as for the real Primates dataset above, except that the tree prior is “Calibrated Yule” for panels b and d of fig. 3. This tree prior was updated to Birth-Death in panel c because it was showing a bias towards younger ages. The number of trees with any ESS under 200 is given in supp. info. S15.

## Code and data availability

The Python statistical analysis, the source data and the intermediate data are archived at Zenodo.org with the DOI: <u>10.5281/zenodo.14000603</u>. The core phylogenetics/bioinformatics library is also available at github.com/DyogenIBENS/Phylorgs (version 0.1.0).

## Acknowledgment

We wish to thank Alexandra Louis for the management of data and software resources used in this work, and Pierre Vincens for computing support.

