## Supplementary information for "Factors influencing the accuracy and precision in dating single gene trees"

2024/10/28

[1]Centre for Anthropobiology and Genomics of Toulouse, CNRS UMR5288, Université Paul Sabatier, Toulouse, France
[2]École Normale Supérieure, PSL Research University, CNRS, Inserm, Institut de Biologie de l'École Normale Supérieure (IBENS), F-75005 Paris, France

**Supplementary information S1** 21 species Primate tree with median branch lengths.

Branch lengths are the median number of substitutions per nucleotide from the 5204 gene trees, obtained from the Beast clock model fit.

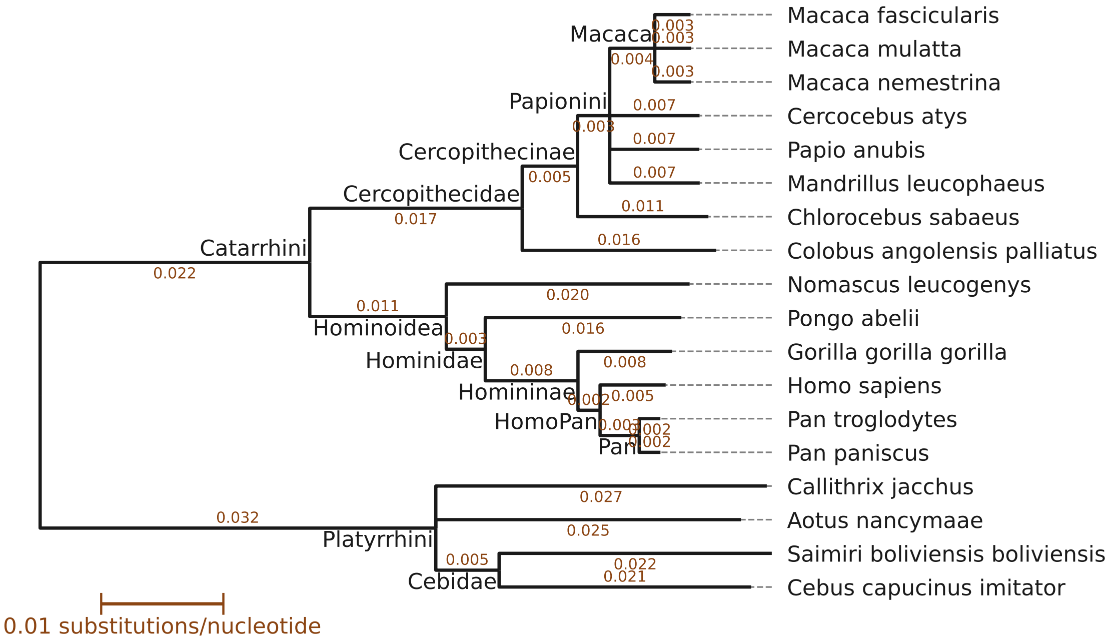

**Supplementary information S2** Complete list of the 56 input features for the regression.

To obtain the final selection of features, the following removal criteria were checked in order:

1. constant features;
2. collinear features upon PCA inspection;
3. features which identify aberrant gene trees (outliers), alongside their corresponding aberrant trees;
4. features not retained by the Lasso fit, or with a coefficient strictly less than 0.01.

Retained variables are highlighted in light green.

| Variable | Description | Removal |
|---|---|---|
| | **Input tree features (reconciled trees)** | |
| aberrant_dists | Number of branches with length > 10000 | (1) |
| rebuilt_topo | Tree topology was forced to fit the species tree | (4) |
| unlike_clock | Root-to-tip length deviation in the Ensembl tree | |
| bootstrap_min | Minimum node bootstrap value (Ensembl) | (4) |
| bootstrap_mean | Mean node bootstrap value (Ensembl) | (2) |
| RF | Gene tree discordance with the species tree (Robinson-Foulds) | |
| | **Alignment features (based on ingroup sequences only)** | |
| ingroup_glob_len | Alignment length | |
| ingroup_mean_GC | Average GC-content of ingroup sequences | (4) |
| ingroup_mean_N | Average proportion of ambiguous nucleotides per sequence | (4) |
| ingroup_mean_gaps | Average proportion of gaps per sequence | (4) |
| ingroup_mean_CpG | Average proportion of CpG dinucleotides per sequence | (2) |
| ingroup_std_len | Standard deviation of sequence lengths | (4) |
| ingroup_std_GC | Std. dev. of the GC-content of ingroup sequences | |
| ingroup_std_N | Std. dev. of the proportion of ambiguous nucleotides per sequence | (2) |
| ingroup_std_gaps | Std. dev. of the proportion of gaps per sequence | (2) |
| ingroup_std_CpG | Std. dev. of the proportion of CpG per sequence | (2) |
| ingroup_nucl_entropy_mean | Mean entropy score across aligned nucleotides | (2) |
| ingroup_nucl_entropy_median | Median entropy score across aligned nucleotides | (4) |
| ingroup_nucl_entropy_std | Std. dev. of the entropy score across aligned nucleotides | (2) |
| ingroup_nucl_parsimony_mean | Mean parsimony score across aligned nucleotides | (2) |
| ingroup_nucl_parsimony_std | Std. dev. of the parsimony score across aligned nucleotides | (4) |
| ingroup_codon_entropy_mean | Mean entropy score across aligned codons | (2) |
| ingroup_codon_entropy_median | Median entropy score across aligned codons | (2) |
| ingroup_codon_entropy_std | Std. dev. of the entropy score across aligned codons | (2) |
| ingroup_codon_parsimony_mean | Mean parsimony score across aligned codons | (2) |
| ingroup_codon_parsimony_median | Median parsimony score across aligned codons | (2) |
| ingroup_codon_parsimony_std | Std. dev. of the parsimony score across aligned codons | (2) |
| | **Output from alignment filtering** | |
| prop_splitseq | proportion of non-overlapping sequences | (3) |
| hmmc_propseqs | Proportion of sequences modified by HmmCleaner | (4) |
| hmmc_max | Maximum proportion of sequence deleted by HmmCleaner | (4) |
| | **Codeml output features** | |
| NsynSites | Amount of synonymous substitutions in the stationary state | |
| brOmega_mean | Average $\omega = dN/dS$ of branches | (4) |
| brOmega_std | Standard deviation of $\omega = dN/dS$ of branches | (4) |
| brOmega_skew | Skew of $\omega = dN/dS$ of branches | (4) |
| ls | Number of aligned sites | (2) |
| | **Beast outputs** | |
| unconverged | Has at least one parameter with ESS < 200 | (3) |
| beast_nsamples | Number of samples in the MCMC chain | (4) |
| treeL_12_med | Total tree length at codon positions 1,2 (posterior median) | (2) |
| treeL_3_med | Total tree length at codon positions 3 | (2) |
| gammaShape_12_med | Shape of gamma site rates at codon positions 1,2 | (4) |
| gammaShape_3_med | Shape of gamma site rates at codon position 3 | |
| kappa_12_med | $\kappa$ at codon positions 1,2 | |
| kappa_3_med | $\kappa$ at codon position 3 | |
| ucldMean_12_med | Beast lognormal clock mean parameter at codon positions 1,2 | (2) |

| | | |
|---|---|---|
| ucldMean_3_med | Beast lognormal clock mean parameter at codon positions 3 | (2) |
| ucldStdev_12_med | Beast lognormal clock standard deviation parameter at codon positions 1,2 | (2) |
| ucldStdev_3_med | Beast lognormal clock standard deviation parameter at codon positions 3 | (2) |
| rate_12_mean_med | Branch average of the Beast rate statistic at codon positions 1,2 | (2) |
| rate_12_var_med | Branch variance of the Beast rate statistic at codon positions 1,2 | (2) |
| rate_3_mean_med | Branch average of the Beast rate statistic at codon positions 3 | (2) |
| rate_3_var_med | Branch variance of the Beast rate statistic at codon positions 3 | (2) |
| | **Rate features** | |
| beastmedian_rate | Branch-average of the Beast rate statistic, in subst/codon/My (posterior median) | |
| beastclockmedian_rate | Beast lognormal clock mean parameter (posterior median) | (2) |
| beastmedian_rate_std | Branch heterogeneity of the rate statistic (sum of std. dev. for all codon positions) | |
| beastclockmedian_rate_std | Branch heterogeneity of the clock rate (sum of the standard deviation of the lognormal clock for all codon positions) | (2) |

**Supplementary information S3** Alternative regression to the one in the main text. Here the only feature for measuring across-branch rate variation is the root-to-tip deviation. The regularization parameter is also $\alpha = 0.02$.

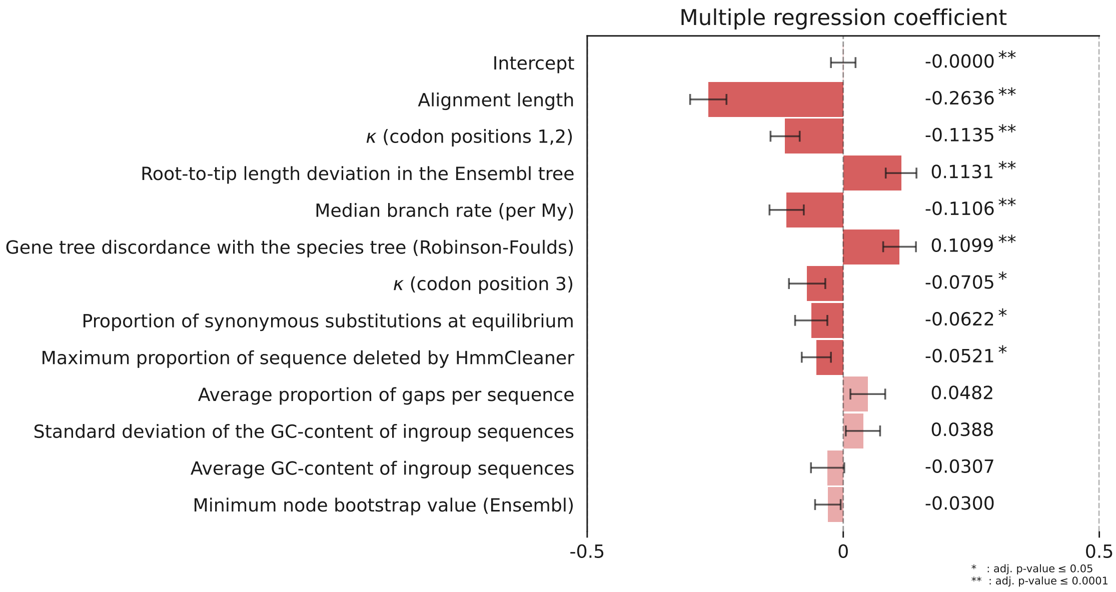

**Supplementary information S4** Enriched human gene annotations from the set with lowest predicted dispersion. Significance threshold 0.05, multiple testing correction by g:SCS (defaults).

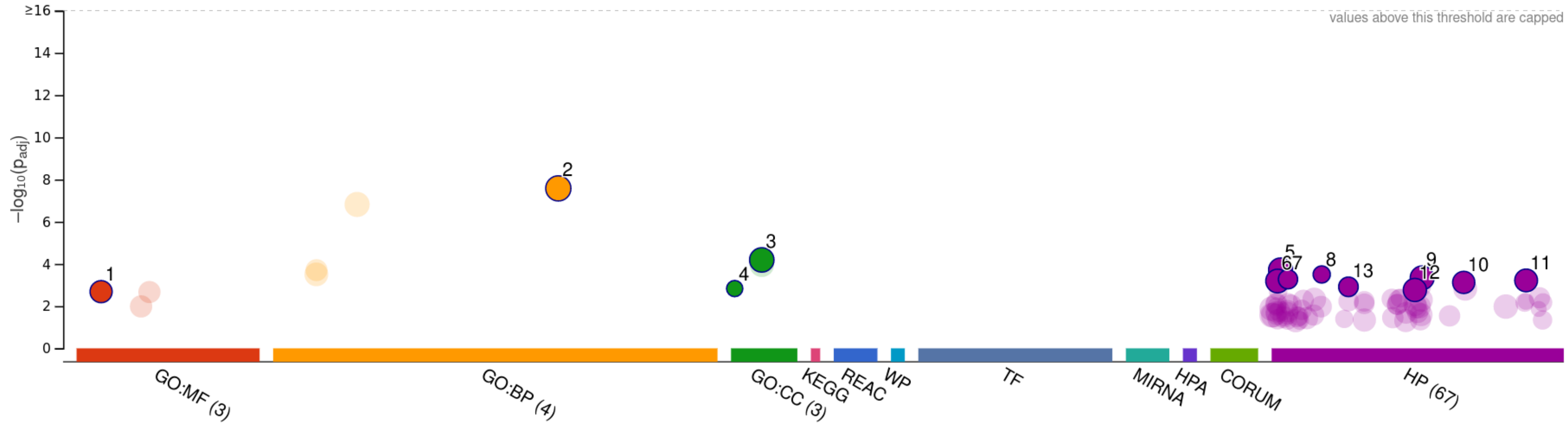


| ID | Source | Term ID | Term Name | $p_{adj}$ (query_1) |
|---|---|---|---|---|
| 1 | GO:MF | GO:0005524 | ATP binding | $2.065\times10^{-3}$ |
| 2 | GO:BP | GO:0071840 | cellular component organization or biogene… | $2.642\times10^{-8}$ |
| 3 | GO:CC | GO:0043228 | non-membrane-bounded organelle | $6.477\times10^{-5}$ |
| 4 | GO:CC | GO:0005604 | basement membrane | $1.475\times10^{-3}$ |
| 5 | HP | HP:0000707 | Abnormality of the nervous system | $1.954\times10^{-4}$ |
| 6 | HP | HP:0000478 | Abnormality of the eye | $6.484\times10^{-4}$ |
| 7 | HP | HP:0001321 | Cerebellar hypoplasia | $5.342\times10^{-4}$ |
| 8 | HP | HP:0004375 | Neoplasm of the nervous system | $3.166\times10^{-4}$ |
| 9 | HP | HP:0012823 | Clinical modifier | $4.589\times10^{-4}$ |
| 10 | HP | HP:0031703 | Abnormal ear morphology | $7.519\times10^{-4}$ |
| 11 | HP | HP:0100022 | Abnormality of movement | $6.059\times10^{-4}$ |
| 12 | HP | HP:0012373 | Abnormal eye physiology | $1.719\times10^{-3}$ |
| 13 | HP | HP:0007360 | Aplasia/Hypoplasia of the cerebellum | $1.221\times10^{-3}$ |

| | |
|---|---|
| **version** | e111_eg58_p18_30541362 |
| **date** | 25/03/2024 09:38:37 |
| **organism** | hsapiens |

g:Profiler

**Supplementary information S5** Enriched human gene annotations from the set of genes with longest alignment. Same parameters as above (defaults).

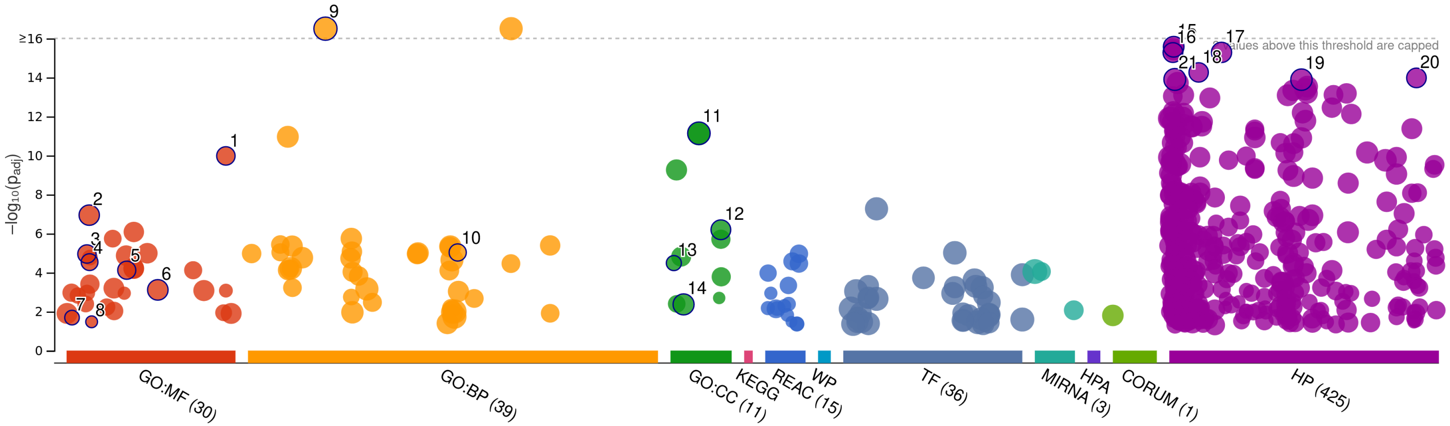


| ID | Source | Term ID | Term Name | $p_{adj}$ (query_1) |
|---|---|---|---|---|
| 1 | GO:MF | GO:0140657 | ATP-dependent activity | $1.052\times10^{-10}$ |
| 2 | GO:MF | GO:0005524 | ATP binding | $1.141\times10^{-7}$ |
| 3 | GO:MF | GO:0005198 | structural molecule activity | $1.119\times10^{-5}$ |
| 4 | GO:MF | GO:0008017 | microtubule binding | $2.924\times10^{-5}$ |
| 5 | GO:MF | GO:0030695 | GTPase regulator activity | $7.597\times10^{-5}$ |
| 6 | GO:MF | GO:0044877 | protein-containing complex binding | $7.736\times10^{-4}$ |
| 7 | GO:MF | GO:0003774 | cytoskeletal motor activity | $2.043\times10^{-2}$ |
| 8 | GO:MF | GO:0008331 | high voltage-gated calcium channel activity | $3.348\times10^{-2}$ |
| 9 | GO:BP | GO:0016043 | cellular component organization | $1.926\times10^{-22}$ |
| 10 | GO:BP | GO:0051056 | regulation of small GTPase mediated signal … | $9.359\times10^{-6}$ |
| 11 | GO:CC | GO:0043232 | intracellular non-membrane-bounded organ… | $7.269\times10^{-12}$ |
| 12 | GO:CC | GO:0099080 | supramolecular complex | $6.406\times10^{-7}$ |
| 13 | GO:CC | GO:0005604 | basement membrane | $3.271\times10^{-5}$ |
| 14 | GO:CC | GO:0030054 | cell junction | $4.296\times10^{-3}$ |
| 15 | HP | HP:0000366 | Abnormality of the nose | $2.663\times10^{-16}$ |
| 16 | HP | HP:0000290 | Abnormality of the forehead | $5.302\times10^{-16}$ |
| 17 | HP | HP:0005105 | Abnormal nasal morphology | $5.186\times10^{-16}$ |
| 18 | HP | HP:0002683 | Abnormal calvaria morphology | $5.518\times10^{-15}$ |
| 19 | HP | HP:0012373 | Abnormal eye physiology | $1.294\times10^{-14}$ |
| 20 | HP | HP:0100886 | Abnormality of globe location | $1.052\times10^{-14}$ |
| 21 | HP | HP:0000478 | Abnormality of the eye | $1.264\times10^{-14}$ |

| | |
|---|---|
| **version** | e111_eg58_p18_30541362 |
| **date** | 31/05/2024 15:40:41 |
| **organism** | hsapiens |

g:Profiler

**Supplementary information S6** Observed distributions of the simulated parameters in the Primates gene trees.

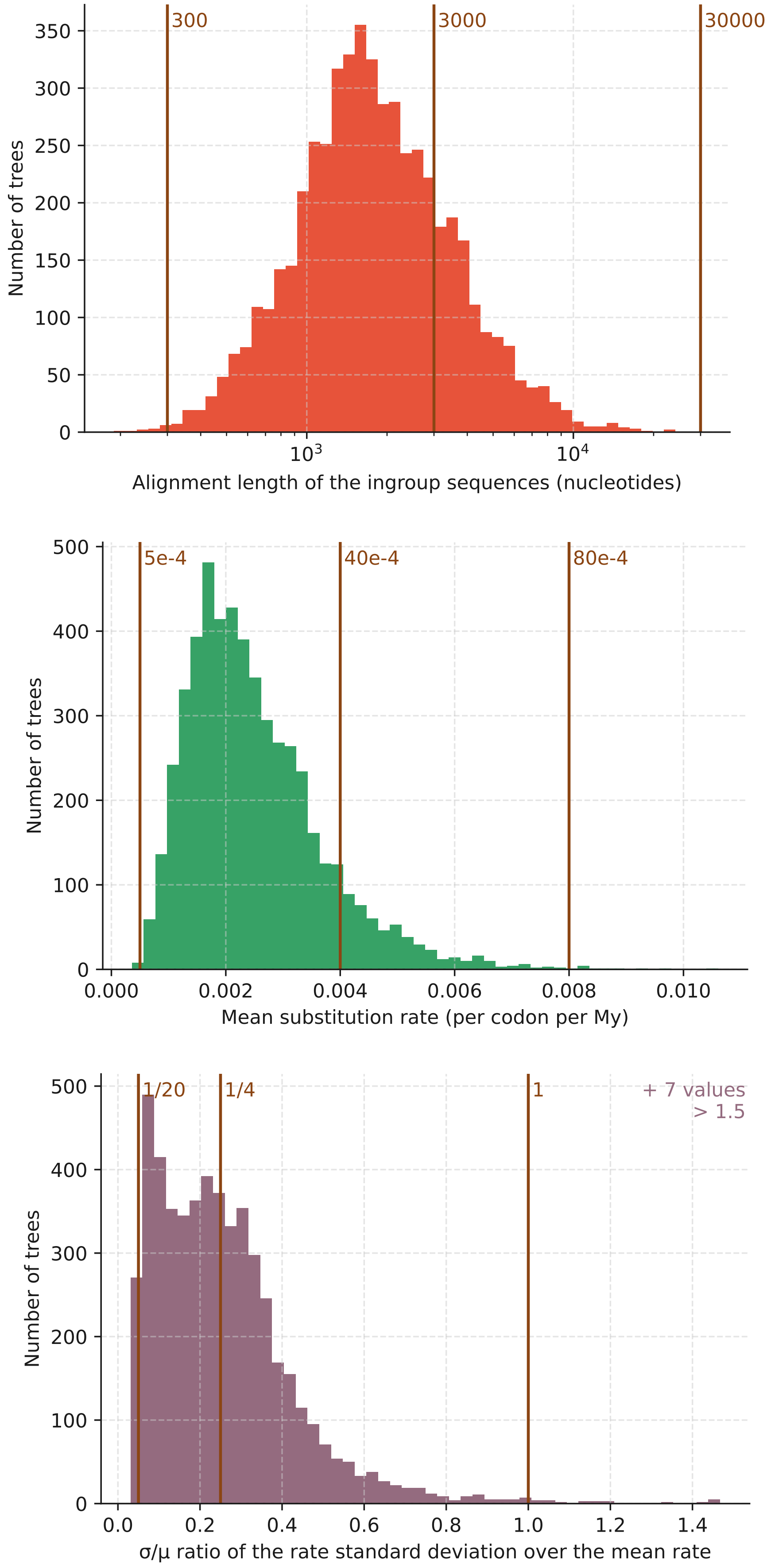

**Supplementary information S7** Sampling from the prior of the Beast model. The underlying tree (black lines) show the dates from TimeTree.

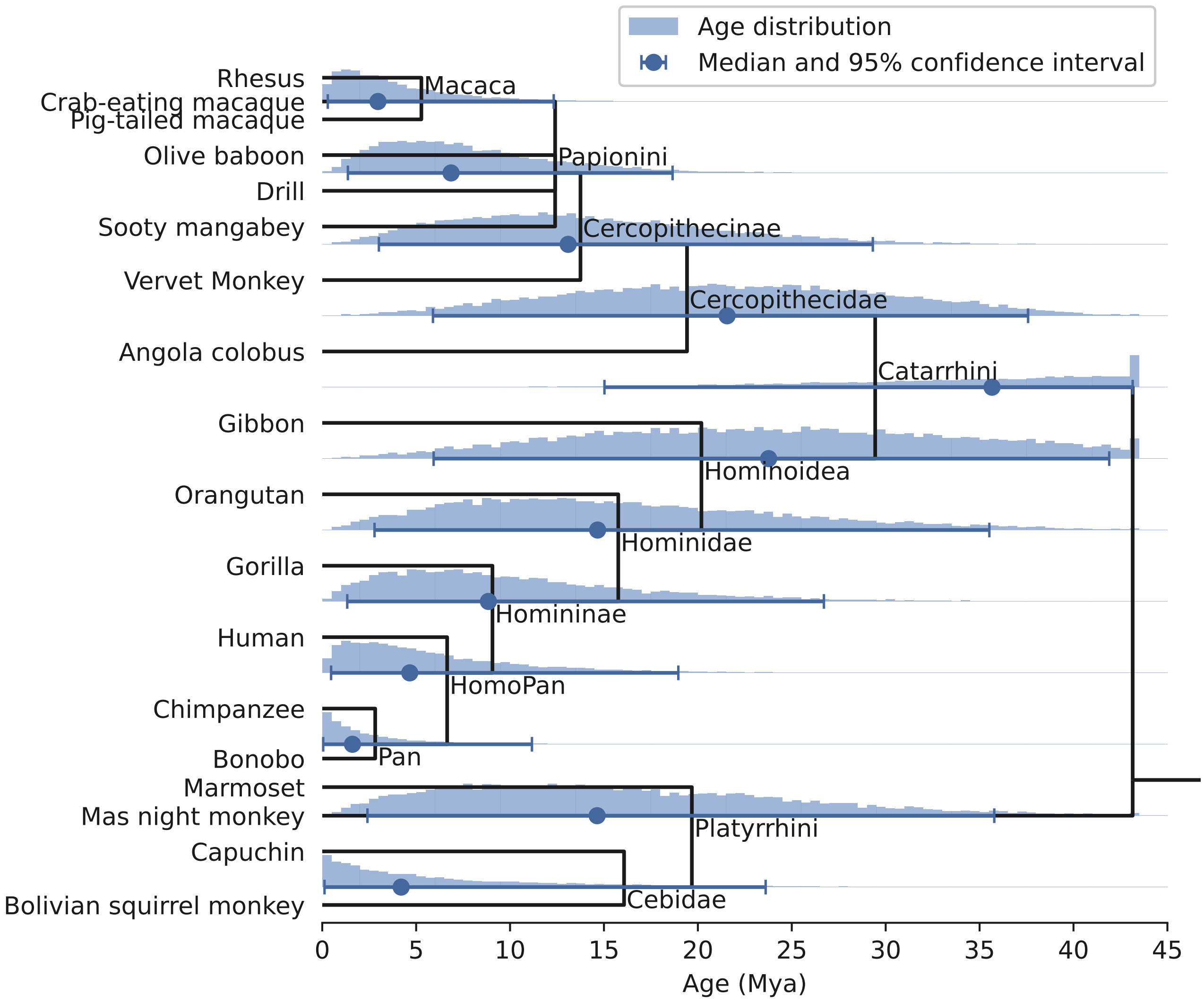

**Supplementary information S8** Species and gene trees preprocessing

The tree of the 99 species from Ensembl Compara 93 is based on the NCBI taxonomy and was edited to resolve polytomies based on multiple sources, including a dated phylogeny from TimeTree (data retrieved Jan. 2019; Kumar et al. 2017) and reference literature (Perelman et al. 2011; Mittermeier et al. 2013; Upham et al. 2019).

We first removed species with low genome assembly quality, as quantified by the $N50$ (size of scaffold in number of genes, such that 50% of genes are contained in larger scaffolds) and the $K70$ (number of the longest scaffolds which contain more than 70% of the genes). Species with $N50 < 30$ genes and $K70 > 400$ scaffolds were removed, including 3 primate species out of 24 (*Rhinopithecus bieti*, *Rhinopithecus roxellana*, *Carlito syrichta*) retaining 21 primate species of which 18 Simiiformes.

Our species tree topology was then applied on the downloaded gene trees from Ensembl, requiring that some nodes be edited (automatically; Peres and Roest Crollius 2015). In the original gene tree forest from Ensembl, 729 subtrees supported by aberrant branch lengths (above 10,000 nucleotide substitutions per site) were removed, and 6962 annotated gene splits (one true gene artefactually annotated as two or more genes fragments) were merged.

**Supplementary information S9** Program versions.

We ran analyses on an Ubuntu 18.04 operating system.

For statistical analyses we used Python 3.8.10 (Jun 22 2022, 20:18:18)

The following Python packages and versions were last used:

| | |
|---|---|
| Numpy | 1.22.3 (GCC 9.4.0) |
| Scipy | 1.8.0 |
| Pandas | 1.4.1 |
| Statsmodels | 0.13.2 |
| Scikit-learn | 1.0.2 |
| Matplotlib | 3.5.3 |
| Seaborn | 0.11.2 |
| Biopython | 1.79 |
| Ete3 | 3.1.2 |
| IPython | 8.4.0 |

**Supplementary information S10** Beast clock model parameters VS rate summary statistics.

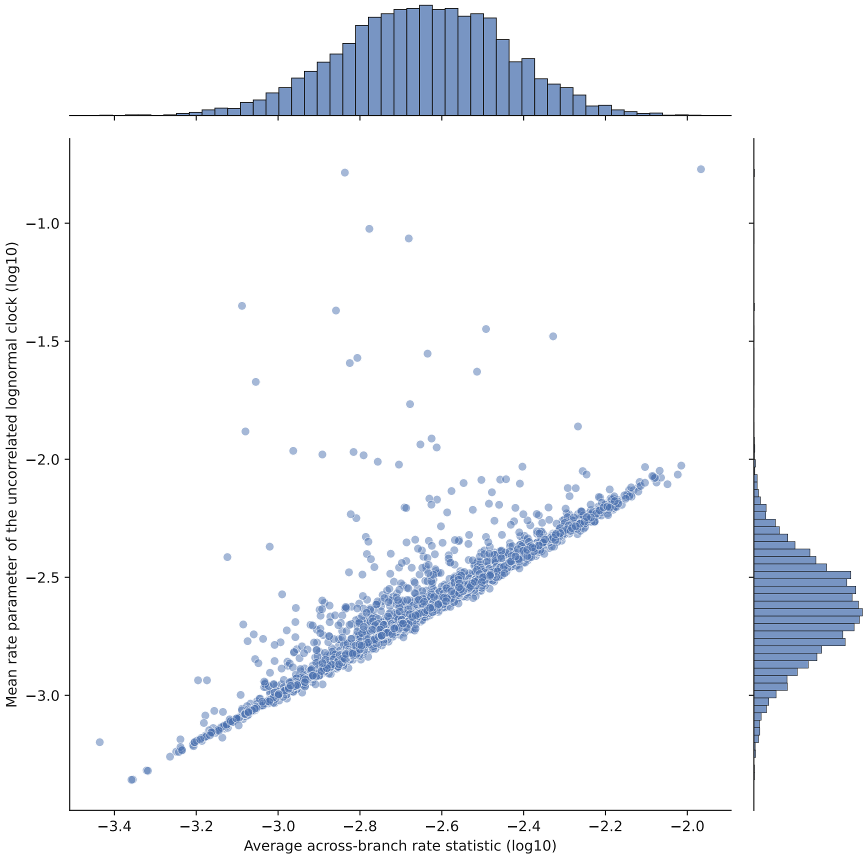


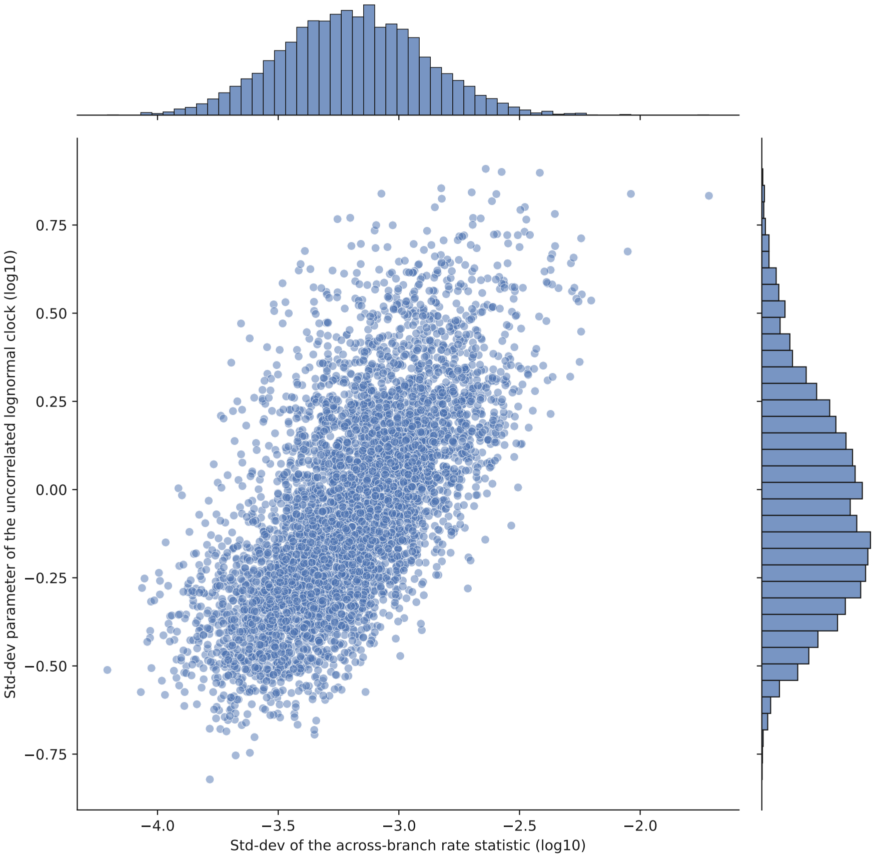

**Supplementary information S11** Feature transformations

Decorrelation operators:

/ division;
− subtraction;
∼ regress and keep the residuals.

| | Transform | Decorrelation | Renaming after decorrelation |
|---|---|---|---|
| rebuilt_topo | `log10(1 + x)` | | |
| unlike_clock | `log10` | | |
| bootstrap_min | `notransform` | | |
| bootstrap_mean | `notransform` | | |
| RF | `sqrt` | | |
| ingroup_glob_len | `log10` | | |
| ingroup_mean_GC | `notransform` | | |
| ingroup_mean_N | `log10(1.2321e-05 + x)` | | |
| ingroup_mean_gaps | `log10(0.00257095 + x)` | | |
| ingroup_mean_CpG | `log10` | / ingroup_mean_GC$^2$ | CpG_odds |
| ingroup_std_len | `log10(3.04024 + x)` | | |
| ingroup_std_GC | `log10` | | |
| ingroup_std_N | `log10(5.08009e-05 + x)` | | |
| ingroup_std_gaps | `sqrt` | | |
| ingroup_std_CpG | `log10` | | |
| ingroup_nucl_entropy_mean | `sqrt` | | |
| ingroup_nucl_entropy_median | `binarize` | | |
| ingroup_nucl_entropy_std | `sqrt` | | |
| ingroup_nucl_parsimony_mean | `log10(0 + x)` | | |
| ingroup_nucl_parsimony_std | `log10(0 + x)` | ∼ ingroup_nucl_parsimony_mean | Ringroup_nucl_parsimony_std |
| ingroup_codon_entropy_mean | `log10` | | |
| ingroup_codon_entropy_median | `binarize` | | |
| ingroup_codon_entropy_std | `sqrt` | | |
| ingroup_codon_parsimony_mean | `log10` | | |
| ingroup_codon_parsimony_median | `binarize` | | |
| ingroup_codon_parsimony_std | `sqrt` | | |
| prop_splitseq | `binarize` | | |
| hmmc_propseqs | `notransform` | | |
| hmmc_max | `log10(0.00676366 + x)` | | |
| treeL_12_med | `-log10(-x)` | | |
| treeL_3_med | `-log10(-x)` | | |
| gammaShape_12_med | `log10` | | |
| gammaShape_3_med | `log10` | | |
| kappa_12_med | `log10` | | |
| kappa_3_med | `log10` | | |
| ucldMean_12_med | `log10` | | |
| ucldMean_3_med | `log10` | | |
| ucldStdev_12_med | `log10` | | |
| ucldStdev_3_med | `log10` | | |
| unconverged | `binarize` | | |
| beast_nsamples | `binarize` (cutoff 10002) | | |
| rate_12_mean_med | `log10` | | |
| rate_12_var_med | `log10` | | |
| rate_3_mean_med | `log10` | | |
| rate_3_var_med | `log10(sqrt(x))` | | |
| NsynSites | `log10` | − ls | RsynSites |
| brOmega_mean | `sqrt` | | |
| brOmega_std | `sqrt` | / brOmega_mean | RbrOmega_std |
| brOmega_skew | `notransform` | | |
| ls | `log10` | | |
| beastmedian_rate | `log10` | | |
| beastclockmedian_rate | `log10` | | |
| beastmedian_rate_std | `log10` | ∼ beastmedian_rate | Rbeastmedian_rate_std |
| beastclockmedian_rate_std | `log10` | | |
| beastclock_rate_extreme | `binarize` | | |
| beastclockmedian_rate_extreme | `binarize` | | |

**Supplementary information S12** Aberrant gene trees (35), removed prior to the regression.

| | Threshold | Number of trees |
|---|---|---|
| Presence of pairs of sequences without overlap (split) | $> 0$ | 16 |
| Presence of an ESS $< 200$ | $> 0$ | 13 |
| Beast lognormal clock mean (posterior median) | 0.03 | 1 |
| Beast lognormal clock mean (posterior mean) | 0.03 | 8 |
| Any of the above | | 35 |

**Supplementary information S13** Detailed outputs and parameters of the regression of per-tree error.

| | coef | std err | $P>\|z\|$ | adj. p-value | [0.025 | 0.975] | Lasso coef | Simple regression coef | Simple regression $R^2$ | Simple regression p-value | Simple regression adj. p-value |
|---|---|---|---|---|---|---|---|---|---|---|---|
| const | -0.0000 | 0.0120 | 1.0000 | nan | -0.0230 | 0.0230 | 0.0000 | -0.0000 | 0.0000 | 1.0000 | nan |
| ingroup_glob_len | -0.3471 | 0.0160 | 0 | 0.0000 | -0.3790 | -0.3160 | -0.3324 | -0.3684 | 0.1357 | 0 | 0 |
| Rbeastmedian_rate_std | 0.3014 | 0.0130 | 0 | 0.0000 | 0.2760 | 0.3270 | 0.2814 | 0.2688 | 0.0723 | 0 | 0 |
| beastmedian_rate | -0.1370 | 0.0150 | 0 | 0.0000 | -0.1670 | -0.1070 | -0.1043 | -0.0920 | 0.0085 | 0 | 0 |
| RF | 0.0770 | 0.0150 | 0 | 0.0000 | 0.0470 | 0.1070 | 0.0773 | 0.3165 | 0.1002 | 0 | 0 |
| kappa_3_med | -0.0764 | 0.0170 | 0 | 0.0001 | -0.1090 | -0.0430 | -0.0184 | -0.1126 | 0.0127 | 0 | 0 |
| RsynSites | -0.0736 | 0.0160 | 0 | 0.0001 | -0.1040 | -0.0430 | -0.0186 | -0.0550 | 0.0030 | 0 | 0.0005 |
| ingroup_std_GC | 0.0584 | 0.0140 | 0 | 0.0008 | 0.0310 | 0.0860 | 0.0398 | 0.1656 | 0.0274 | 0 | 0 |
| unlike_clock | 0.0477 | 0.0130 | 0 | 0.0063 | 0.0220 | 0.0730 | 0.0454 | 0.2278 | 0.0519 | 0 | 0 |
| gammaShape_3_med | -0.0470 | 0.0150 | 0 | 0.0372 | -0.0760 | -0.0180 | -0.0115 | -0.0042 | 0.0000 | 0.7884 | 1 |
| kappa_12_med | -0.0427 | 0.0140 | 0 | 0.0645 | -0.0700 | -0.0150 | -0.0505 | -0.2255 | 0.0509 | 0 | 0 |

| | Simple regression [0.025 | Simple regression 0.975] | VIFs | transform | transformed mean | transformed std | decorr |
|---|---|---|---|---|---|---|---|
| const | -0.0273 | 0.0273 | 1.0000 | `nan` | nan | nan | `nan` |
| ingroup_glob_len | -0.3939 | -0.3430 | 1.8128 | `log10` | 3.2501 | 0.2893 | |
| Rbeastmedian_rate_std | 0.2400 | 0.2976 | 1.2069 | `log10` | -3.3346 | 0.3854 | ∼ `beastmedian_rate` |
| beastmedian_rate | -0.1208 | -0.0632 | 1.4776 | `log10` | -2.6491 | 0.2009 | |
| RF | 0.2888 | 0.3443 | 1.6035 | `sqrt` | 1.6409 | 1.0329 | |
| kappa_3_med | -0.1399 | -0.0853 | 1.9512 | `log10` | 0.9177 | 0.1632 | |
| RsynSites | -0.0801 | -0.0299 | 1.7783 | `log10` | 2.7077 | 0.3008 | `NsynSites − ls` |
| ingroup_std_GC | 0.1380 | 0.1931 | 1.3596 | `log10` | -2.3732 | 0.2742 | |
| unlike_clock | 0.1996 | 0.2561 | 1.1800 | `log10` | -0.7710 | 0.2383 | |
| gammaShape_3_med | -0.0346 | 0.0262 | 1.4072 | `log10` | -0.0600 | 0.1827 | |
| kappa_12_med | -0.2556 | -0.1955 | 1.2601 | `log10` | 0.6129 | 0.1456 | |

Column descriptions:

**coef** coefficient from the multiple regression (OLS fit);
**std err** standard error of the previous coefficient;
**P>|z|** p-value of the coefficient being different from zero (yellow background means significant at level 0.05);
**adj. p-value** p-value adjusted by a Bonferroni correction;
[**0.025**, **0.975**] 95% confidence interval upper/lower limits;
**Lasso coef** coefficient of the lasso regression used to select features;
**Simple regression coef, $\mathbf{R^2}$, p-value, ...** for comparison with the multiple regression, simple regressions (i.e., testing one variable at a time, with intercept) were computed;
**VIF** Variance Inflation Factor. This metric characterizes the collinearity added by a variable. Values above 10 should be avoided.
**transform** the function applied to the raw variable prior to regressing;
**transformed mean, std** mean and standard deviation of the variable post-transform;
**decorr** how, and against which variable a variable was "decorrelated". "∼" indicates that the residuals from a simple regression were kept.

**Supplementary information S14** Observed distribution of $\kappa$ (transitions/transversions) in the Primates gene trees.

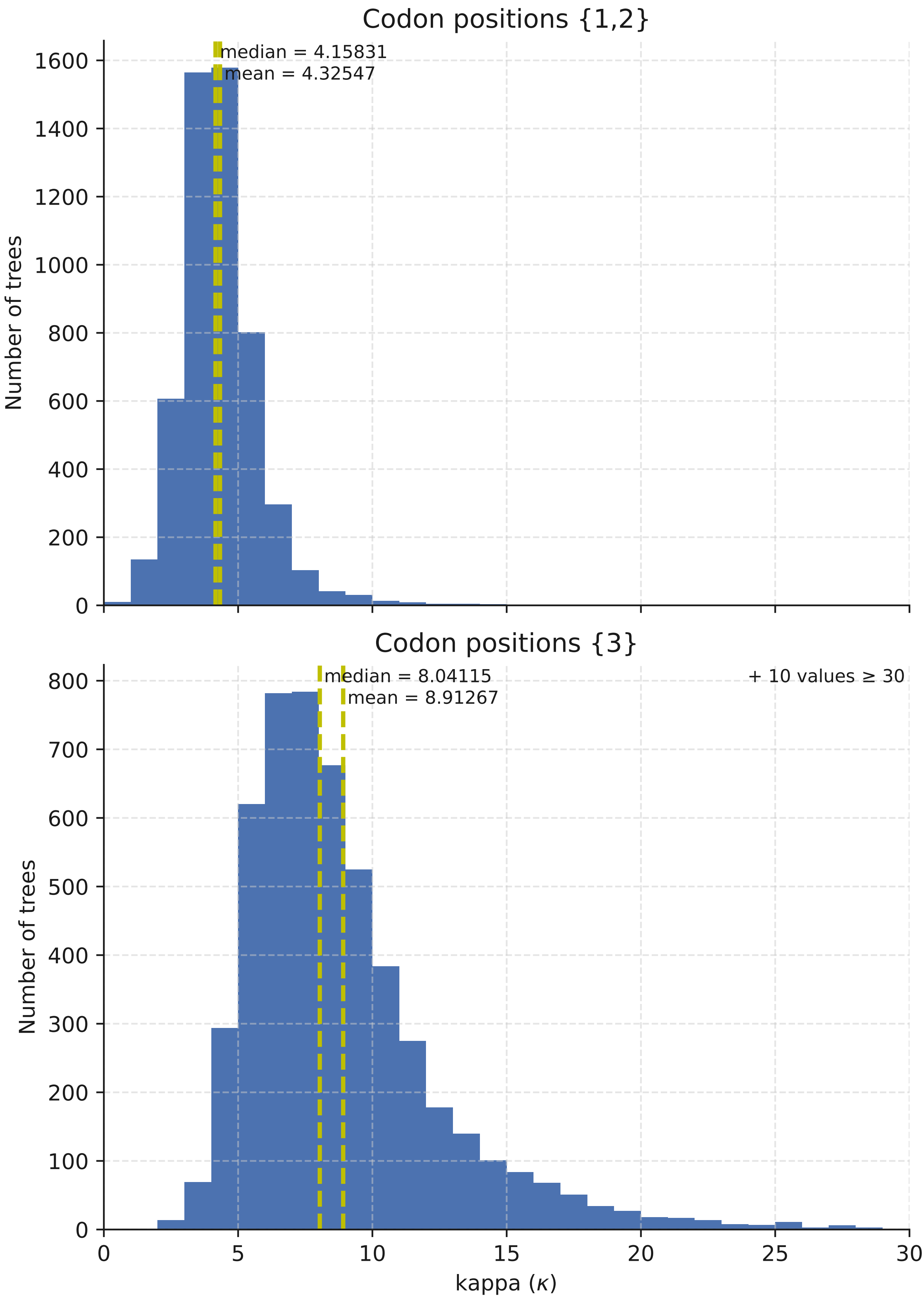

**Supplementary information S15** Number of trees having one or more variable with ESS<200

| **Dataset** | | | **Number of trees with ESS<200** |
|---|---|---|---|
| **Empirical** | | | |
| 5205 Primates gene trees | | | 13 |
| **Simulated (500 replicates each)** | | | |
| Alignment length | mean rate $\mu$ | rate heterogeneity $\sigma/\mu$ | |
| 300 | $40 \times 10^{-4}$ | 1/4 | 1 |
| 3000 | $40 \times 10^{-4}$ | 1/4 | 1 |
| 30000 | $40 \times 10^{-4}$ | 1/4 | 182 |
| 3000 | $5 \times 10^{-4}$ | 1/4 | 6 |
| 3000 | $80 \times 10^{-4}$ | 1/4 | 0 |
| 3000 | $40 \times 10^{-4}$ | 1/20 | 6 |
| 3000 | $40 \times 10^{-4}$ | 1 | 51 |
